# Relative precision of the sibship and LD methods for estimating effective population size with genomics-scale datasets

**DOI:** 10.1101/2021.06.15.448561

**Authors:** Robin S. Waples

## Abstract

Computer simulations were used to compare relative precision of two widely-used single-sample methods for estimating effective population size (*N_e_*)—the sibship method and the linkage-disequilibrium (LD) method. Emphasis is on performance when thousands of gene loci are used, which now can easily be achieved even for non-model species. Results show that unless *N_e_* is very small, if at least 500-2000 diallelic loci are used, precision of the LD method is higher than the maximum possible precision for the sibship method, which occurs when all sibling relationships have been correctly identified. Results also show that when precision is high for both methods, their estimates of effective population size are high and positively correlated, which limits additional gains in precision that might be obtained by combining information from the two estimators.

## Introduction

Effective population size (*N_e_*) is an important parameter in evolutionary biology, but direct calculation requires detailed demographic information that is difficult to obtain for natural populations. For this reason, genetic methods to estimate *N_e_* have been used for the past half century. For most of this time period, the vast majority of genetically-based *N_e_* estimates used the temporal method, which quantifies the rate of genetic drift over time and requires at least two samples separated by one or more generations (Krimbas and Tsakas 1971; Nei and Tajima 1981; Waples 1989; Wang 2001; Wang and Whitlock 2003). This changed dramatically in the late 2000s following development of two methods that require only a single sample: the bias-adjusted method based on linkage disequilibrium (LD; Waples 2006; Waples and Do 2008) and the sibship method of Wang (2009). Within just a few years, new publications using these single-sample methods far exceeded those using various versions of the temporal method (Palstra and Fraser 2012).

Performance of the LD and sipship methods has been extensively evaluated with simulated and empirical data (e.g., Wang 2009, 2016; Waples and Do 2010; Waples et al. 2014; Gilbert and Whitlock 2015; Ackerman et al. 2017). However, most of these evaluations have used no more than a few dozen microsatellite markers or their equivalent in diallelic, single-nucleotide-polymorphism (SNP) loci. The study using the largest number of markers to directly compare the two methods was by Wang (2016), who simulated 1000 diallelic (’SNP’) loci and measured root-mean-squared-error (RMSE, which reflects both bias and precision) as a function of genome size, measured in Morgans. Wang found that RMSE was higher for the LD method with small genomes (likely reflecting bias due to physical linkage) but that RMSE was essentially the same for the two methods when genome size was larger than about 20 Morgans. For perspective, total genome size is estimated to be 36.1 Morgans in humans (Kong et al. 2002) and 32.5 in cattle (Arias et al. 2009).

Wang’s (2016) simulations using 1000 SNPs considered only a single scenario—true *N_e_* = 50, with estimates based on sampling 50 offspring—and therefore only scratch the surface of the vast parameter space potentially of interest to researchers. This is an important data gap, as recent technological advances now make it relatively easy to generate large genomics-scale datasets (10^3^−10^6^ or more SNPs), even for non-model species. Two factors have made such assessments challenging in the past and have contributed to this data gap. First, Wang’s method for inferring sibling relationships (implemented in the software *Colony*; Jones and Wang 2010) is computationally demanding because it jointly considers groups of individuals in computing close-kin likelihoods, rather than treating each pair of individuals independently (as, for example, is done by *ML-relate*, Kalinowski et al. 2006). This makes it difficult for other researchers to evaluate the sibship method with large samples of individuals or loci. The second challenging factor is that the large numbers of markers used in genomics-scale datasets all must be packaged into a relatively small number of chromosomes. Loci close together on the same chromosome do not assort independently and do not provide independent information about evolutionary processes, and this reduces precision in large datasets, but by an amount that is difficult to quantify. Furthermore, the LD method depends on averaging the LD signal across many pairs of loci, and in theory precision increases quickly for large-scale studies because the number of pairwise comparisons of *L* loci is *L*(*L*−1)/2 ≈ *L*^2^/2. But the many locus pairs all share overlapping subsets of the same *L* loci, and this creates another type of pseudoreplication that reduces precision.

Fortunately, it is possible to overcome both of these limitations to provide a direct comparison of performance of the sibship and LD methods with large genomics datasets. A recent study (Waples et al. 2020) has shown that precision of the LD method can be accurately predicted based on four covariates: *N_e_*, sample size, number of loci, and number of chromosomes (a measure of genome size). Somewhat surprisingly, this study found that the major factor limiting precision of the LD method in large genomics datasets is not physical linkage itself but rather the lack of independence caused by averaging the LD signal across many overlapping pairs of loci. Waples et al. (2020) found that smaller genomes do reduce precision more; however, this effect attenuates rapidly, such that precision for species with 16 chromosomes was only slightly reduced compared to those with 64 chromosomes, and that results for 64 chromosomes were largely indistinguishable from results for scenarios that simulated unlinked loci. This means that unlinked loci can be used to model precision of the LD method, with a minor adjustment to account for effects of genome size.

Implementing Wang’s sibship method in an ambitious simulation study that uses large numbers of loci remains very challenging. However, there is a simple workaround that takes advantage of the fact that precision in the sibship method depends on the number of sibling matches that are found, just as precision in traditional mark-recapture methods for estimating abundance depends on the number of tag recoveries in the second sample (Otis et al. 1978). Wang’s method is categorical in the sense that each pair of offspring either produce a sibling match or not, so the number of matches is always an integer. This means that once the number of genetic markers used is sufficient to reliably identify the true pedigree of sampled individuals, precision cannot be increased by addition of more loci. Thus, while it is still difficult to accurately measure absolute precision of the sibship method, it is relatively easy to identify an upper limit to precision, which can be measured by keeping track of the true pedigree and assuming that all kin inferences are made without error. In contrast, the LD method depends on mean *r*^2^ (the squared correlation of alleles at different gene loci) averaged across many pairs of loci, which is a continuous variable whose variance declines as more loci are used, but at an increasingly slower rate in large genetic datasets (as recently quantified by Waples et al. 2020).

Here, simulations are used to quantify precision of the LD estimator for a number of realistic scenarios (ranges of *N_e_*, sample size, and number of loci) and compare that with the maximum possible precision for the sibship method. In addition, the correlation structure of the two estimators was evaluated to help understand the relative benefits of developing a combined estimator using information from both methods.

## Methods

Simulations were conducted in R (R Core Team 2021) using code provided in the Appendix. Modeled populations followed the original Wright-Fisher model: monecious diploids with random mating (including random selfing), discrete-generations, and a constant size of *N* ideal individuals. Under these conditions, *N_e_* = *N* (on average). To minimize extra variance caused by random variation in realized *N_e_* each generation (which has variance ≈*N*/2; Waples and Faulkner 2009), vectors of offspring number per parent were randomly cycled through and only those that produced realized *N_e_* within 0.5% of the target value were used.

In each replicate, 10 generations of burnin were run to allow mean *r*^2^ to reach a dynamic equilibrium, and then data were collected for the next 20 generations. Fifty replicates were run for each scenario, which produced a total of 1000 replicate *N_e_* estimates for each of the methods. Four different effective sizes were simulated [*N_e_* = 50, 200, 1000, 5000], and for each effective size three different sample sizes of individuals were analyzed, chosen to produce roughly equivalent ranges of precision (see Figure 1). Each year in each replicate, genetic data for 5000 diallelic (SNP) loci were generated, and mean *r*^2^ was calculated for the full complement of loci, as well as subsets of 100-2500 loci. Calculation of *r*^2^ and the bias adjustment for estimating *N_e_* followed Waples (2006), so the resulting estimates were the same as produced in the program LDNe (Waples and Do 2008). The initial simulations included >5000 loci but loci with minor allele frequency < 0.05 were removed, and those remaining were randomly trimmed to 5000 to ensure consistent numbers for analysis.

**Figure 1.**
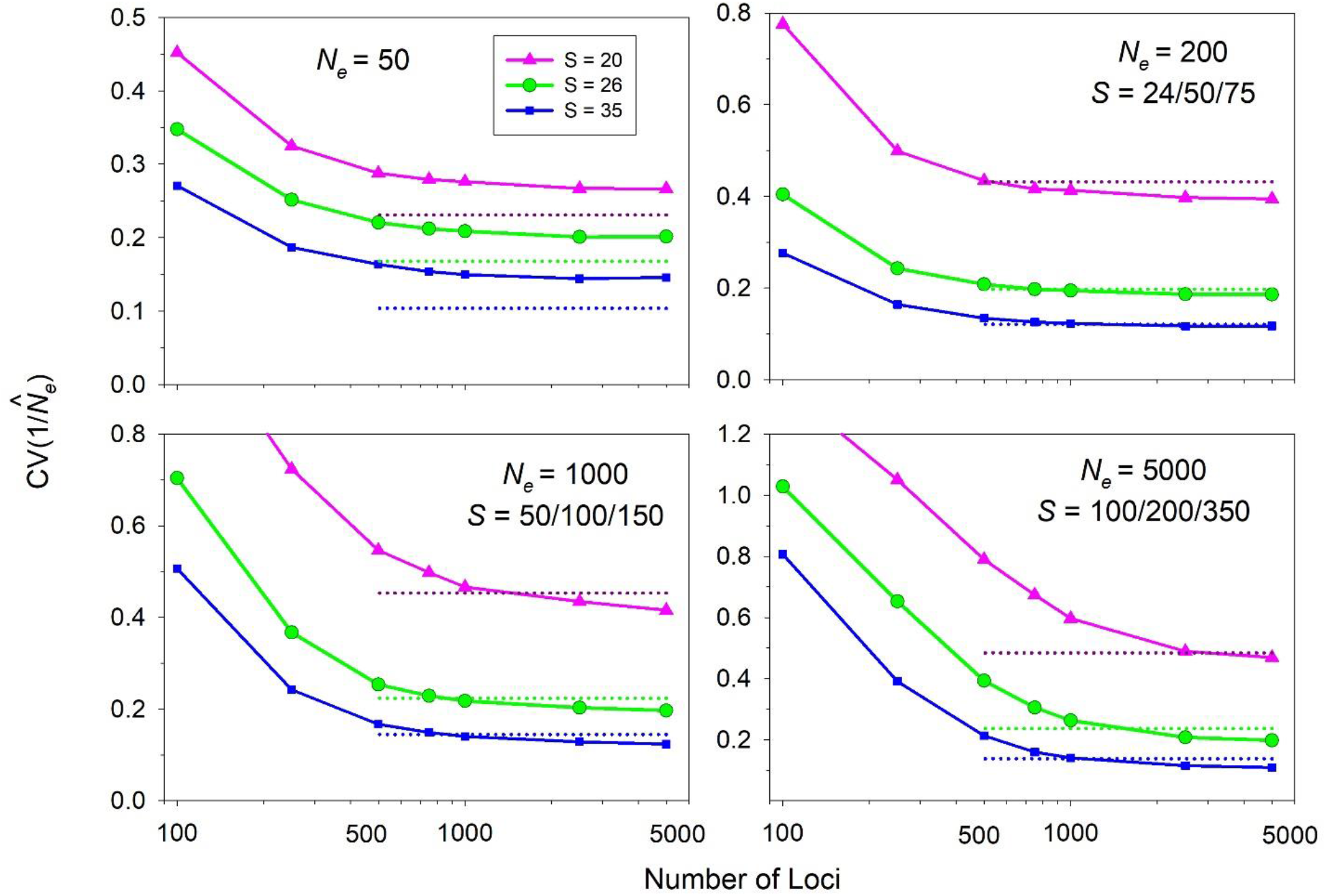
Comparison of CVs of 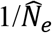 for the LD method and minimum possible CVs for the sibship method, under the assumption that all sibling relationships are identified without error. For each effective size, results are shown for three different sample sizes of offspring (*S*); in each case the pink lines and symbols depict results for the smallest sample size, green lines and symbols depict results for intermediate sample sizes, and blue lines and symbols depict results for the largest sample sizes. Dotted lines, color coded to the respective sample sizes, show minimum CVs for the sibship method. Note the log scale on the *X* axis and the different linear scales on the *Y* axis.

In each year of data collection, parents for all sampled offspring were recorded, and these data were used to identify full and half siblings. Following Wang (2009), sibling frequencies across all pairwise comparisons of individuals are denoted by *Q_FSP_* and *Q_HSP_*, with the latter representing all half siblings (maternal and paternal combined). These frequencies can then be used to estimate effective size as (Wang 2009):

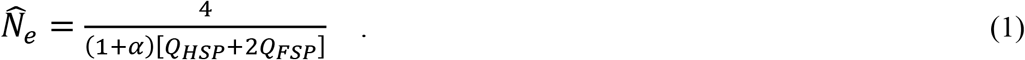

In Equation 1, the term α accounts for effects of selfing, which in our model occurred with probability 1/*N_e_*. Following Wang (2016), α was calculated as α = (1/*N_e_*)/(2 – 1/*N_e_*), which is very close to 1/(2*N_e_*). Our simulations used *N_e_* = [50,200,1000,5000], for which α computes as [0.01, 0.005, 0.001, and 0.0005]—in all cases representing a small correction that had little effect on the results.

To quantify precision, the coefficient of variation (CV) was computed across replicate estimates of effective size for both methods. Because of the inverse relationship between 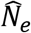 and sibling frequencies (Equation 1), even if the observed number of sibling matches is approximately normally distributed (as will often be the case), 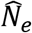 will not be; instead, 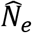is skewed high and is infinitely large when no recoveries are found. A similar issue applies to the LD method, which can even return a negative estimate if observed mean *r*^2^ is less than the value expected to arise from sampling error alone. For this reason, it is common for evaluations of precision of effective size estimators to focus on the distribution of 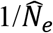 rather than 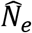(e.g., Wang 2001; 2009). Accordingly, here we report values for the CV of 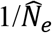. Any infinite estimates for the sibship method or negative estimates for the LD method were converted to large positive numbers (99999) for computing the CVs.

Although Waples et al. (2020) found that most of the pseudoreplication in LD analyses arises from overlapping pairs of loci used to compute mean *r*^2^—and this effect is captured by simulating unlinked loci as in the present study—there is also a modest effect of genome size. This genome-size effect was accounted for using R code provided by Waples et al. (2020) that allows users to predict how much different covariates affect the variance of mean *r*^2^. First, we calculated an adjustment factor that was the ratio of the predicted CV of *r*^2^ for a species with 20 chromosomes (a typical number for many organisms) to the predicted CV for unlinked loci. Predicted CVs are higher for smaller genomes, so these adjustment factors were ≥1; values ranged from 1 to 1.2 depending on the scenario, and were generally higher for smaller effective sizes and sample sizes (see Table A1). Then, the raw CV values obtained in the simulations were multiplied by these adjustment factors to obtain adjusted CV values for mean *r*^2^, and these adjusted CVs were used in all subsequent analyses. This adjusted the empirical CVs for the LD method to what would be expected for an organism with 20 chromosomes.

If more than one method is used on the same data to estimate effective size, a combined estimate can have better performance than either method does by itself. Whether this is the case or not depends on two factors: a) relative biases associated with the methods, and b) whether the methods provide independent or correlated information about *N_e_*. A considerable body of data shows that the sibship and LD methods are largely unbiased when model assumptions are met, but the correlation structure of the two estimators has not been evaluated. To quantify this, for each combination of *N_e_*, sample size, and number of loci, we computed the Pearson product-moment correlation between 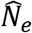 for the two methods, using the two vectors of 1000 replicate estimates.

## Results

Averaged across 1000 replicate samples for each scenario, both methods produced essentially unbiased estimates of effective population size (Figure A1). Results comparing empirical CVs of 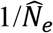 for the LD method with minimum CVs for the sibship method showed a qualitative difference between small and moderate to large *N_e_* (Figure 1). As expected (and as found by Waples et al. 2020), the CV for the LD method dropped consistently as more loci were used, but at an increasingly slower rate. In contrast, the minimum possible CV for the sibship method is independent of the number of loci, because it assumes the genetic data are sufficient to correctly identify all siblings. For *N_e_* = 50, the CV for the LD method never dropped as low as the minimum CV for the sibship method, even using 5000 loci, and Waples et al. (2020) found that increasing the number of loci beyond that number did little to further reduce the variance of mean *r*^2^ when *N_e_* was only 50. In contrast, for all the larger *N_e_* values that were evaluated, the CV for the LD method dropped below the minimum CV for the sibship method by the time 500-2000 loci were used, and the gap between the two methods continued to widen as more loci were added. With 5000 loci, the CV for the LD method was 3-21% lower (depending on sample size) than the minimum CV for the sibship method when *N_e_* was 5000, and reductions for *N_e_* = 1000 and 200 were 8-14% and 3-9%, respectively.

Correlations between estimated *N_e_* for the two methods are affected in a complex way by true *N_e_* and sample sizes of loci and individuals (Figure 2). Most correlation were positive, some very strongly so (*r* > 0.9), and in general the strength of the correlation increased with *N_e_*, sample size, and number of loci. However, for *N_e_* = 200 the strongest correlations were found for the intermediate sample size, and for *N_e_* = 50 the strongest correlations were found for the smallest sample size.

**Figure 2.**
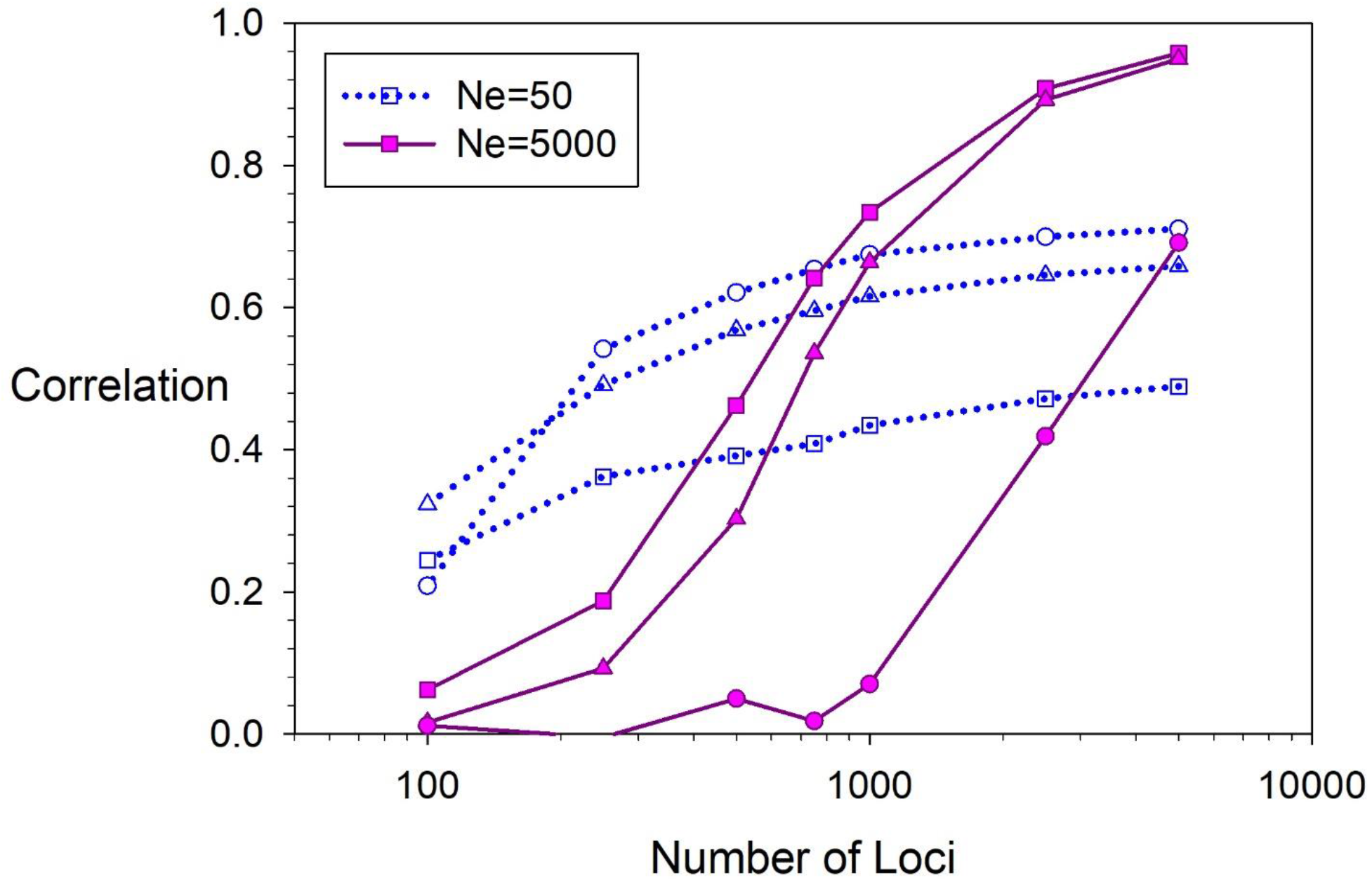
Correlations between 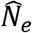 for the LD and sibship methods for scenarios with true *N_e_* = 50 (dotted blue lines and open symbols) and *N_e_* = 5000 (solid pink lines and filled symbols). For each effective size, square symbols represent the largest sample size, triangles intermediate sample size, and circles the smallest sample size.

## Discussion

This study produced two new results that are of direct relevance to practical applications. First, using only relatively modest numbers of loci (~500-2000) that now can be easily generated even for non-model species, precision of the LD method for estimating effective population size can equal or exceed the maximum possible precision for the sibship method. The differences in precision are greatest for large effective sizes, which is a useful result because the drift signal for large *N_e_* is so small that it can be difficult to distinguish from sampling error (Waples 2016a; Marandel et al. 2019). Furthermore, as demonstrated by Waples et al. (2020), when *N_e_* is large, precision of the LD method continues to increase well beyond the 5000 maximum loci simulated in this study, which means that in large genomics datasets, CVs can be reduced below those reported here.

An exception to this pattern of relative precision occurs with very small *N_e_*. Results presented here show that a transition occurs somewhere in the *N_e_* range of 50-200, below which point precision of the LD method no longer can reach the maximum possible level of precision for the sibship method. Presumably this occurs because the rate of decay of LD as a function of distance in centimorgans between a pair of SNPs is inversely related to *N_e_* (Sved and Feldman 1973; Weir and Hill 1980), which means that for the same genome size the effects of physical linkage are stronger when effective size is small. Whether this means that, when large numbers of loci are available but *N_e_* is small, precision of the sibship method will be higher than that of the LD method cannot be determined from this study. Tightly-linked loci produce redundant information for pedigree reconstruction just as they do for the LD method, and reliably distinguishing half siblings from unrelated or more distantly-related pairs can be very challenging, even with large amounts of data. Therefore, a more robust comparison of the two methods for small *N_e_* and large numbers of loci will have to await future evaluations that explicitly model uncertainty in the identification of siblings.

The second novel result reported here is that, when precision of the sibship and LD methods are both high, their estimates of *N_e_* are highly and positively correlated. Presumably this occurs because both methods are sensitive to the same signal of inbreeding and identity by descent that is generated by the inbreeding effective size. This result is reassuring, as it shows that two very different methods converge on the same answer as the amount of data increases. On the other hand, these strong correlations are inconvenient because they indicate that, for large genetic datasets, there is only limited potential to further increase precision by computing a combined estimate using results from both methods. A general approach to combining results from two methods would be to calculate a weighted harmonic mean of the two 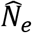 estimates, with weights being inversely proportional to variances of the two estimators (Waples and Do 2010). Waples (2016b) showed that the LD and standard temporal methods produce estimates that are largely uncorrelated, and that when they have roughly equal precision a combined estimate has substantially higher precision than either method alone. It is clear from Figure 2 that estimates from the LD and sibship methods are more strongly correlated than those of the LD and temporal methods. Results presented here provide the information necessary to produce a combined sibship-LD estimate, but only using the ‘best-case’, most optimistic version of the variance associated with 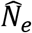 for the sibship method. As shown in Figure 2, correlations between the two estimators decline with reductions in the number of loci used, but the declines in correlations shown in Figure 2 are entirely due to reductions in precision of the LD method. In reality, precision of the sibship method also must decline as fewer loci are used, so the real correlations between estimates provided by the two methods must drop off more sharply with reductions in the number of loci than is shown in Figure 2. Refining these patterns of correlation between the two methods is another topic for future research. To date, most studies that have estimated *N_e_* with both of these methods have used relatively few loci, in which case the estimators are largely uncorrelated and combining results can potentially produce a substantial increase in precision.

Evaluations here have focused on precision, but obtaining reliable *N_e_* estimates from the LD method with large genetic datasets also requires one to deal with potential biases caused by physical linkage. Two general options are available for dealing with this issue. If one does not have detailed linkage information but can estimate genome size or the number of chromosomes (e.g., using data for a related species), the bias adjustment proposed by Waples et al. (2016) can be used to obtain an essentially unbiased estimate of *N_e_*. Alternatively, if the loci can be assigned to chromosomes or linkage groups, mean *r*^2^ can be computed using only pairs of loci on different chromosomes (an option that is implemented in V2 of NeEstimator; Do et al. 2014). Conveniently, the variance of mean *r*^2^ does not depend on whether locus pairs on the same chromosome are used or not (Waples et al. 2020), so choice of which option to use should not affect conclusions about precision.

This study did not generate any new empirical data except by computer simulation. Code to conduct the simulations is available at the end of Supporting Information.

## Supporting Information

**Figure A1.**
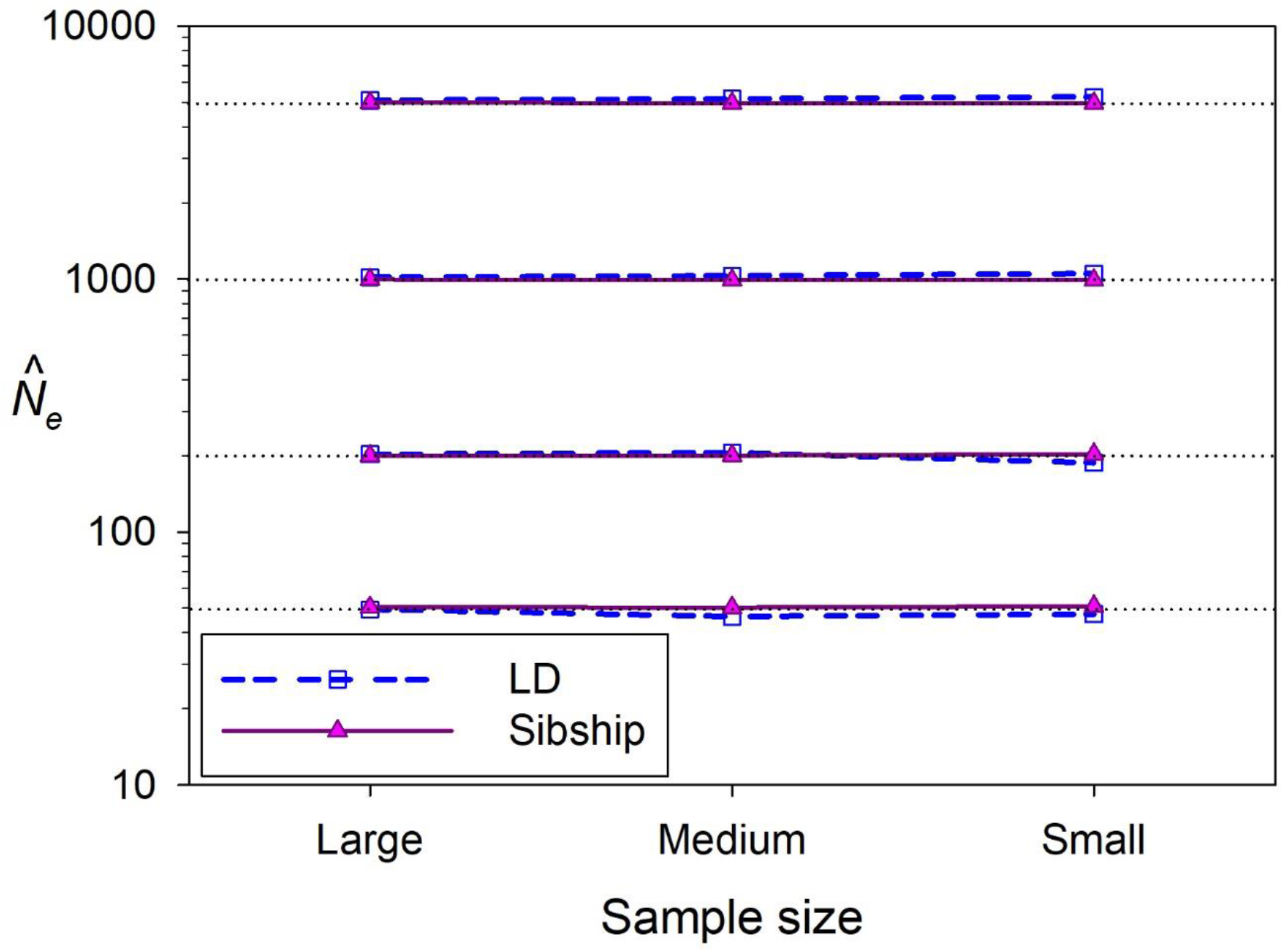
Comparison of 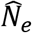 for the LD method (dashed blue lines and open symbols) and the sibship method (solid pink lines and filled symbols) for four different values of true *N_e_* (dotted black lines). Each symbol represents the harmonic mean 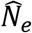 across all replicates for large, medium, and small sample sizes.

**Table A1.** Adjustment factors to account for effects of physical linkage (finite genome size) in the LD analyses. Each value in the table reflects the ratio of the predicted CV for a species with 20 chromosomes of size 50×10^6^ DNA base pairs to the predicted CV that would apply to unlinked loci (as were modeled in this study). Predicted CVs were based on results from Waples et al. (2020). Code from that study was modified for the current analyses and can be found at the end of this supplement. These adjustment factors were used to adjust the raw CVs for the LD method upwards to account for effects of linkage.

| Ne | Loci | Sample size |  |  |
| --- | --- | --- | --- | --- |
|  |  | 20 | 26 | 35 |
| Ne 50 | 100 | 1.216 | 1.188 | 1.175 |
|  | 250 | 1.210 | 1.178 | 1.155 |
|  | 500 | 1.203 | 1.175 | 1.147 |
|  | 750 | 1.200 | 1.174 | 1.146 |
|  | 1000 | 1.198 | 1.174 | 1.146 |
|  | 2500 | 1.194 | 1.177 | 1.153 |
|  | 5000 | 1.192 | 1.179 | 1.159 |
| Ne 200 |  | 24 | 50 | 75 |
|  | 100 | 1.194 | 1.167 | 1.159 |
|  | 250 | 1.196 | 1.164 | 1.149 |
|  | 500 | 1.178 | 1.154 | 1.136 |
|  | 750 | 1.168 | 1.149 | 1.131 |
|  | 1000 | 1.162 | 1.146 | 1.130 |
|  | 2500 | 1.149 | 1.141 | 1.130 |
|  | 5000 | 1.145 | 1.139 | 1.131 |
| Ne 1000 |  | 50 | 100 | 150 |
|  | 100 | 1.095 | 1.089 | 1.087 |
|  | 250 | 1.121 | 1.103 | 1.096 |
|  | 500 | 1.125 | 1.103 | 1.092 |
|  | 750 | 1.118 | 1.099 | 1.087 |
|  | 1000 | 1.112 | 1.096 | 1.085 |
|  | 2500 | 1.095 | 1.088 | 1.081 |
|  | 5000 | 1.089 | 1.085 | 1.081 |
| Ne 5000 |  | 100 | 200 | 350 |
|  | 100 | 0.999 | 0.999 | 0.998 |
|  | 250 | 1.004 | 1.000 | 0.998 |
|  | 500 | 1.018 | 1.005 | 0.999 |
|  | 750 | 1.029 | 1.011 | 1.000 |
|  | 1000 | 1.036 | 1.016 | 1.003 |
|  | 2500 | 1.040 | 1.028 | 1.015 |
|  | 5000 | 1.035 | 1.030 | 1.023 |

### R code for conducting the simulations (which used R version 4.0.4)

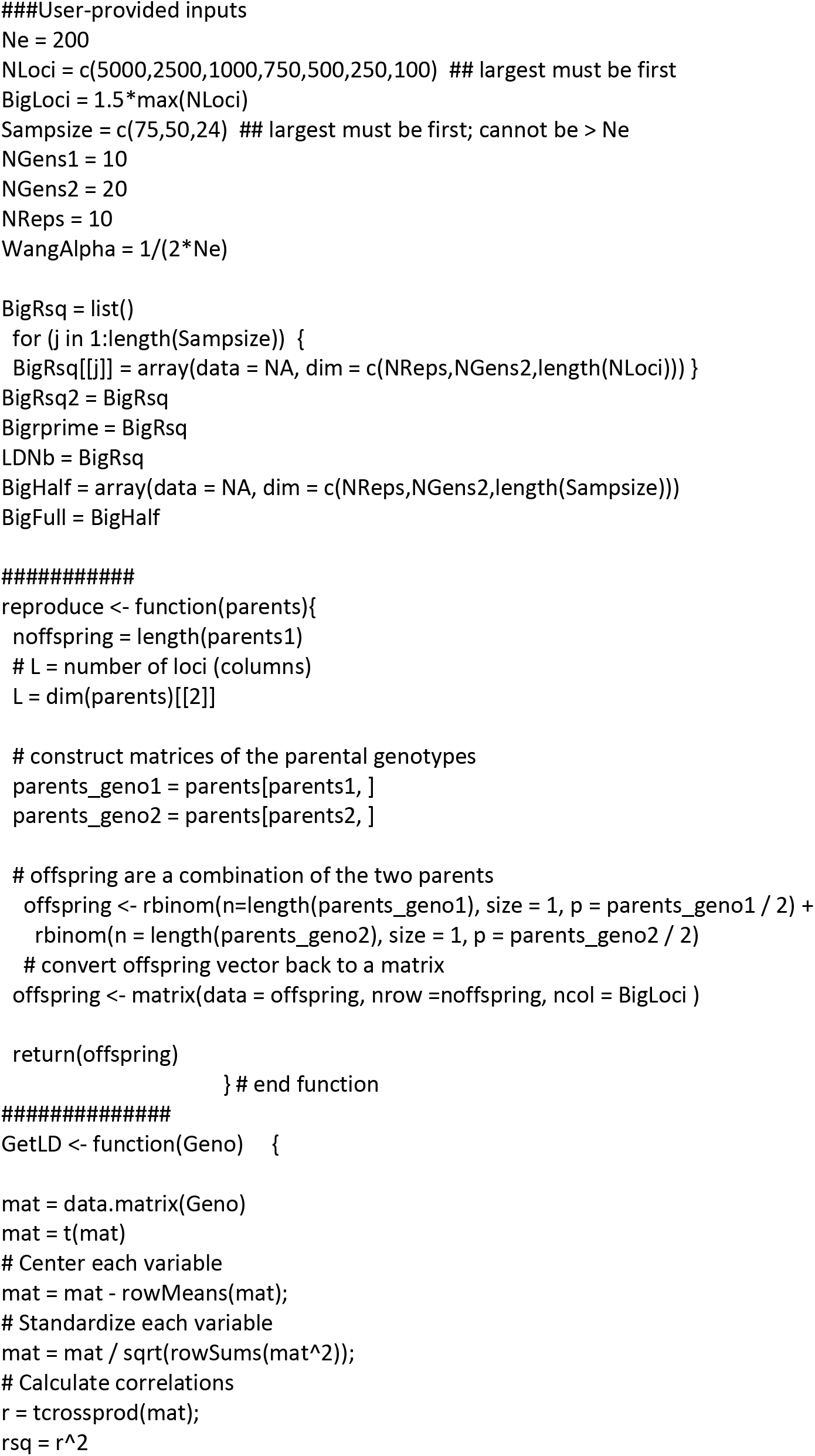
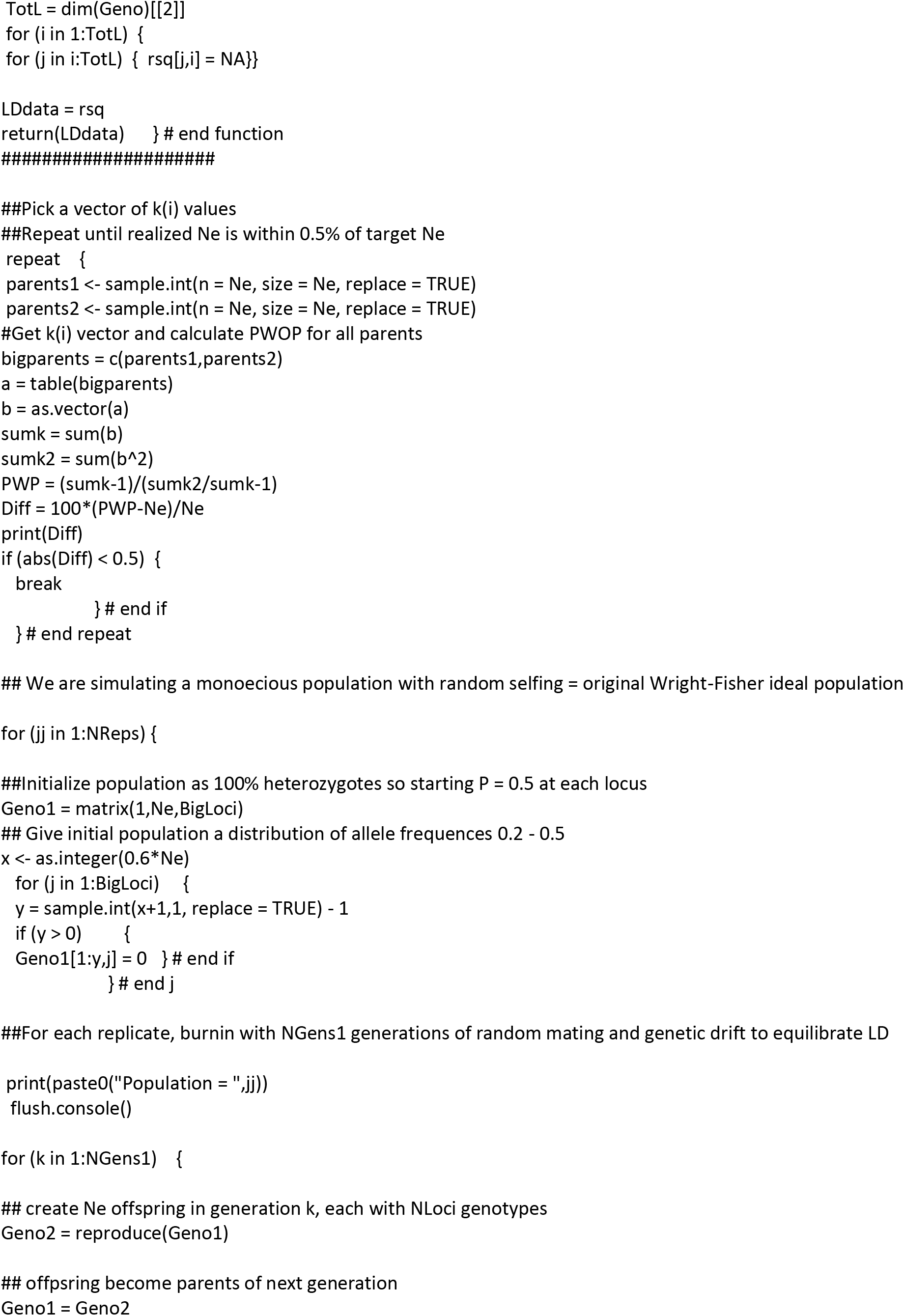
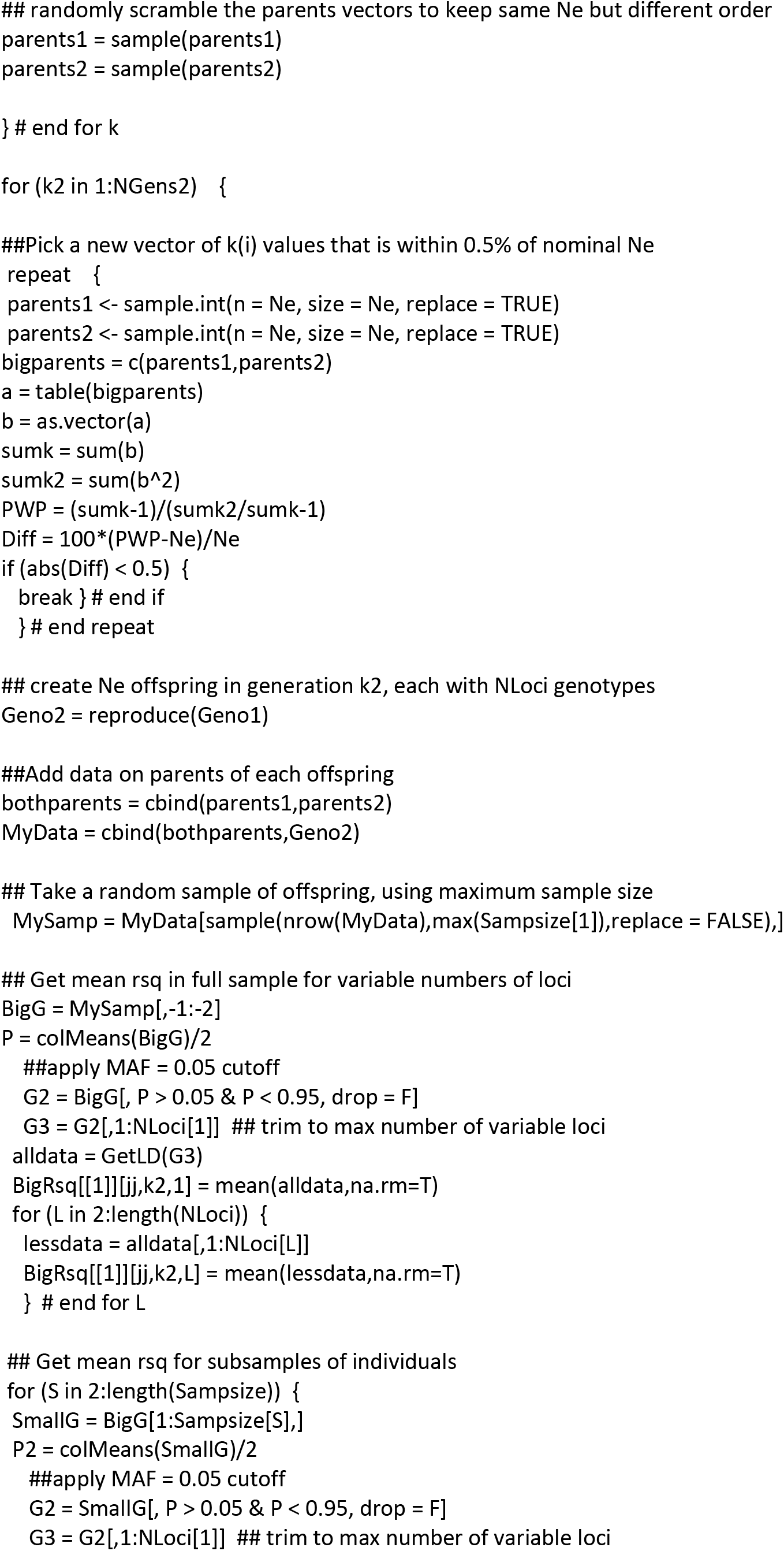
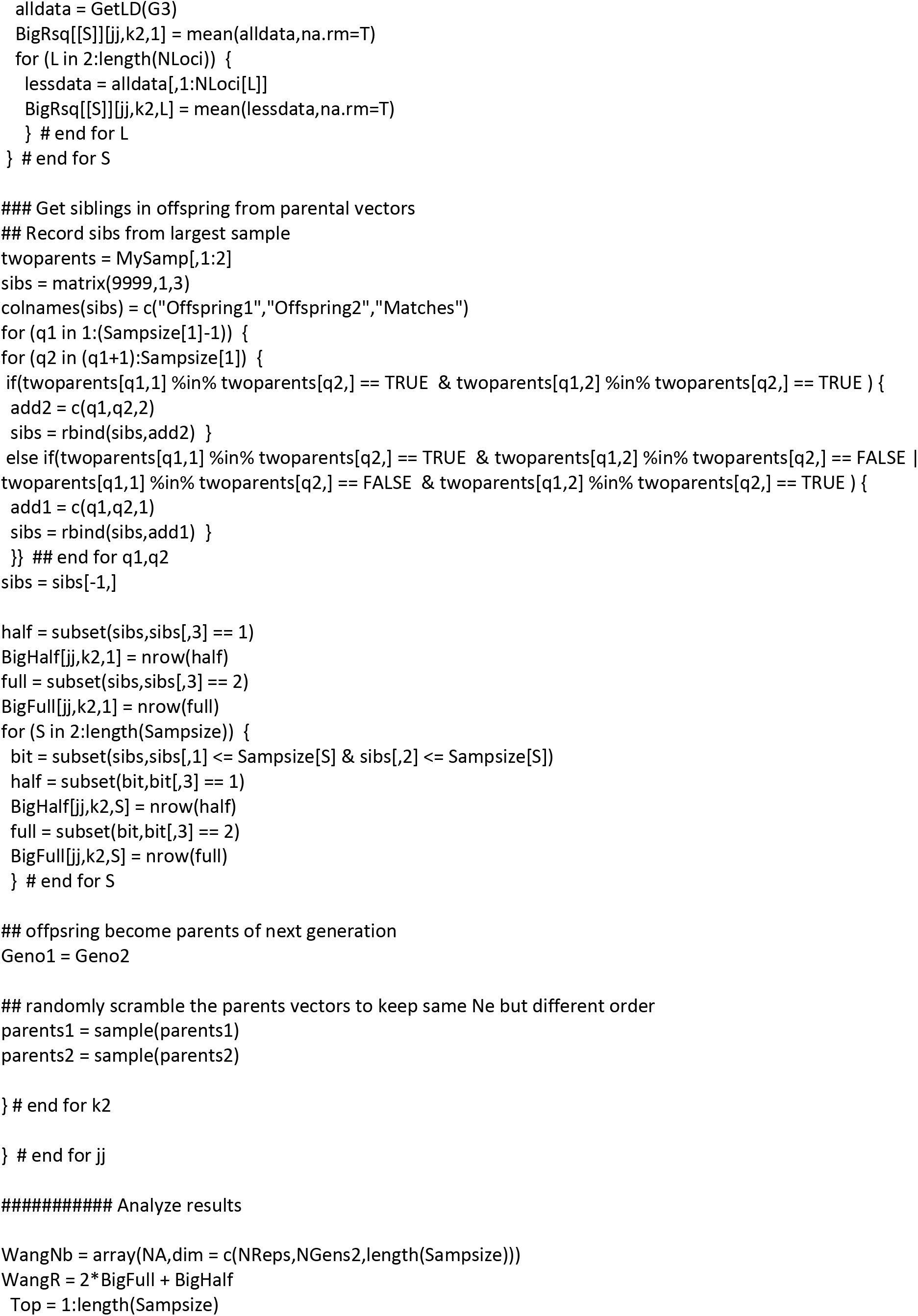
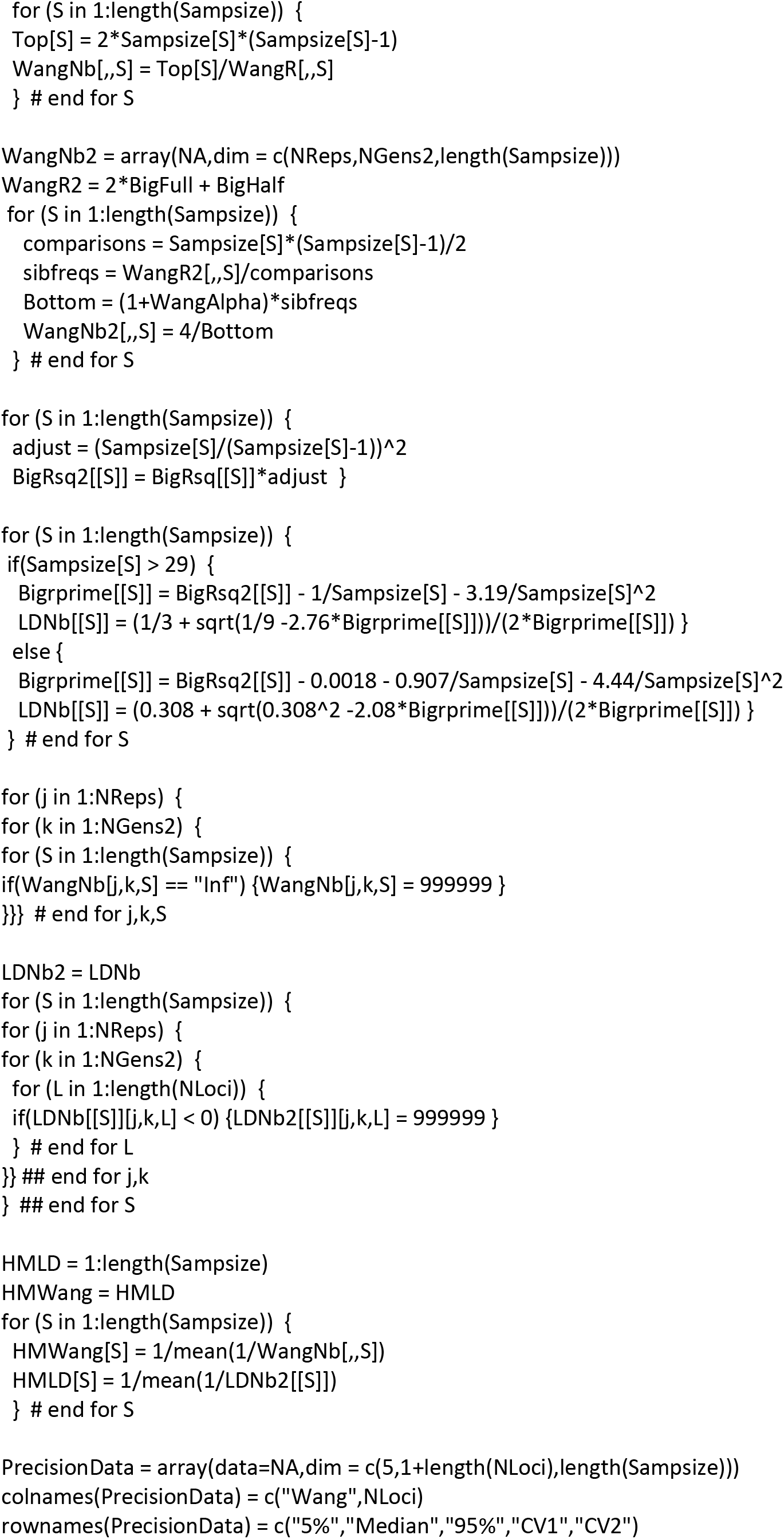
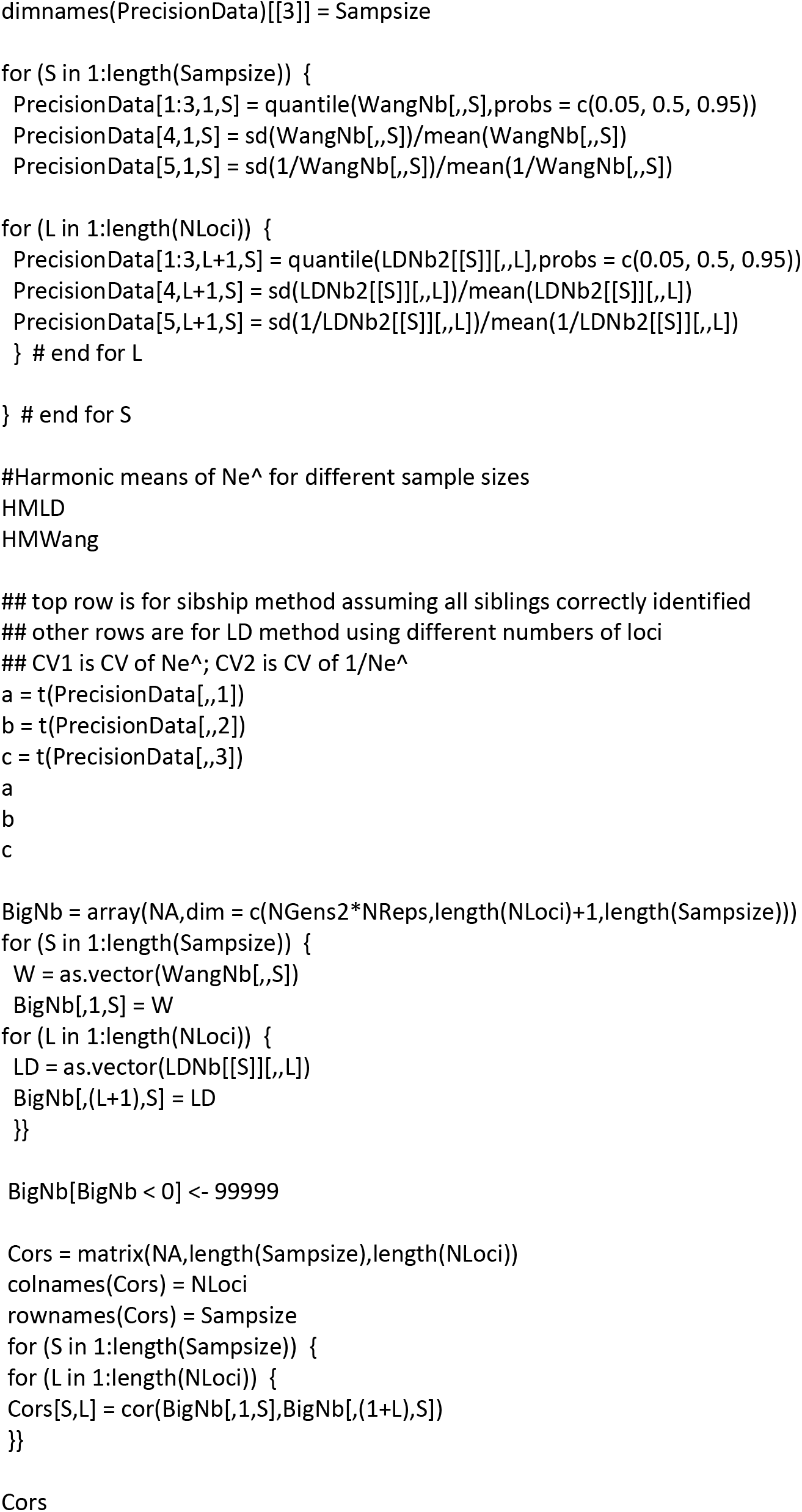

### R code for calculating the adjustment to CV (1/Ne^) for LD method to account for linkage

This code is modified from that used by Waples et al. 2020, Pseudoreplication in genomics-scale datasets. bioRxiv (doi: https://doi.org/10.1101/2020.11.12.380410). The code used in that paper predicts nprime for LD as a function of covariates [Ne,Chr,S,NLoci]. nprime is the effective number of pairwise comparisons of diallelic loci. Here the code is modified to predict the ratio of CV(mean r^2) for an organism with 20 chromosomes to CV(mean r^2) when all loci are unlinked

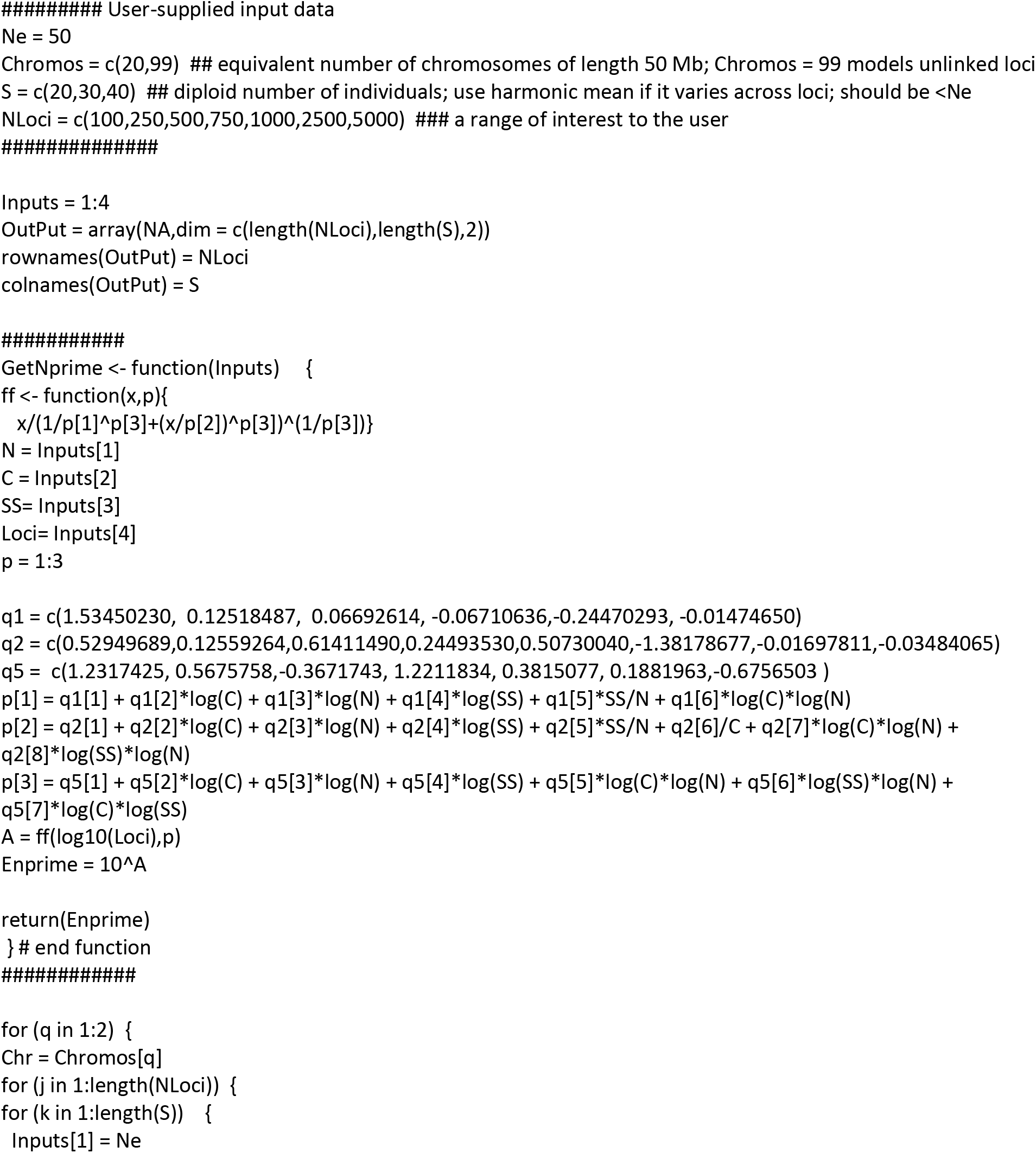
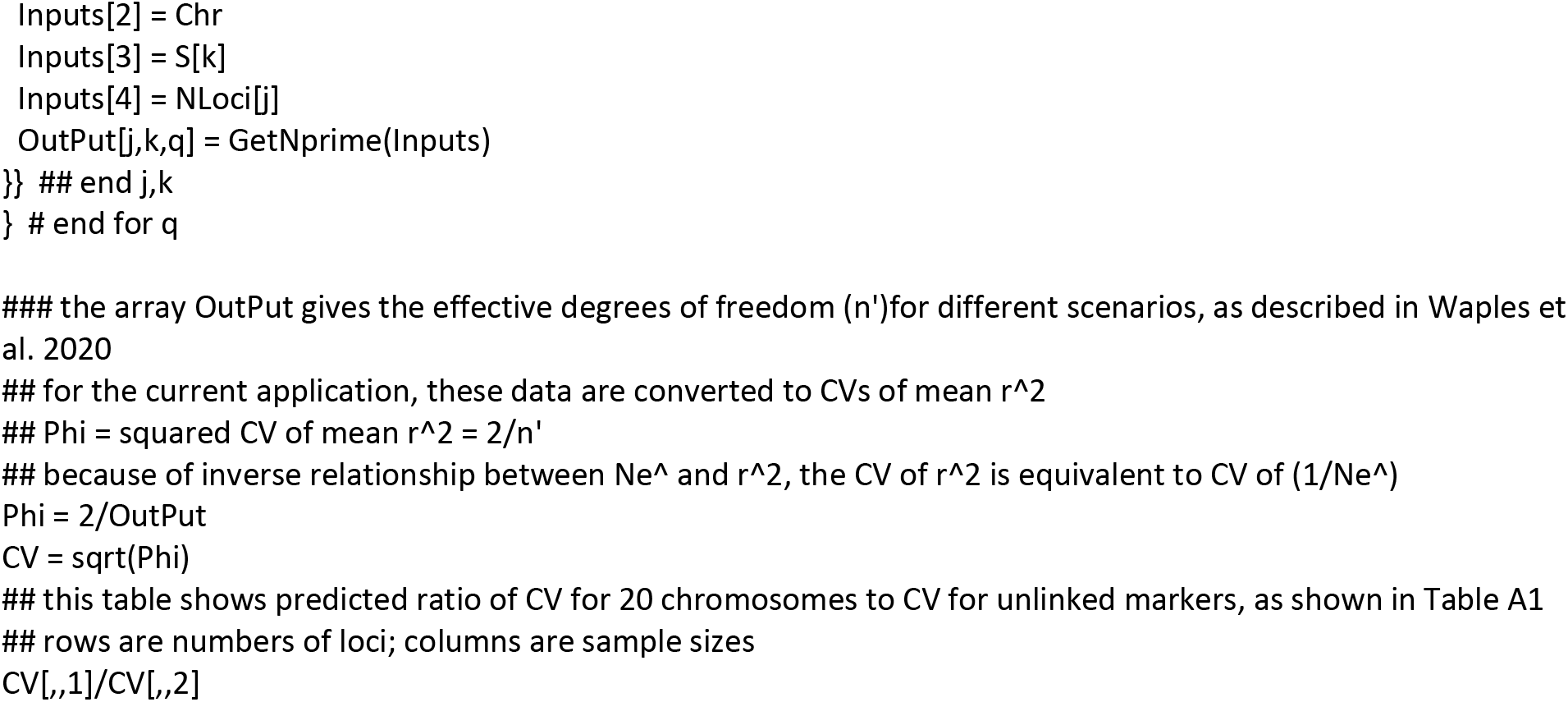

## References

Ackerman, MW, BK Hand, RK Waples, G Luikart, RS Waples, C Steele, BA Garner, J McCane, and M Campbell. 2017. Effective number of breeders from sibship reconstruction: empirical evaluations using hatchery steelhead. Evolutionary Applications 10:146–160.

Arias, J.A., Keehan, M., Fisher, P., Coppieters, W. and Spelman, R., 2009. A high density linkage map of the bovine genome. BMC genetics, 10(1), pp.1–12.

Do, C., R.S. Waples, D. Peel, G.M. Macbeth, B.J. Tillet, and J.R. Ovenden. 2014. NeEstimator V2: re-implementation of software for the estimation of contemporary effective population size (Ne) from genetic data. Molecular Ecology Resources 14:209–214

Gilbert KJ, Whitlock MC (2015) Evaluating methods for estimating local effective population size with and without migration. Evolution, 69, 2154–2166.

Jones, O.R. and Wang, J., 2010. COLONY: a program for parentage and sibship inference from multilocus genotype data. Molecular ecology resources, 10(3), pp.551–555.

Kalinowski ST, Wagner AP, Taper ML (2006) ML-RELATE: a computer program for maximum likelihood estimation of relatedness and relationship. Molecular Ecology Notes, 6, 576–579.

Kong, A., Gudbjartsson, D.F., Sainz, J., Jonsdottir, G.M., Gudjonsson, S.A., Richardsson, B., Sigurdardottir, S., Barnard, J., Hallbeck, B., Masson, G. and Shlien, A., 2002. A high-resolution recombination map of the human genome. Nature genetics, 31(3), pp.241–247.

Nei M, Tajima F (1981) Genetic drift and estimation of effective population size. Genetics 98:625–640.

Krimbas, C.B. and Tsakas, S., 1971. The genetics of Dacus oleae. V. Changes of esterase polymorphism in a natural population following insecticide control—selection or drift?. Evolution, 25(3):454–460.

Marandel F, Lorance P, Berthelé O, Trenkel VM, Waples RS, Lamy JB. 2019. Estimating effective population size of large fish populations, is it feasible? Fish and Fisheries 20:189–198.

Otis, D.L., Burnham, K.P., White, G.C. and Anderson, D.R., 1978. Statistical inference from capture data on closed animal populations. Wildlife monographs, (62), pp.3–135.

Palstra, F.P., and D.J. Fraser, 2012 Effective/census population size ratio estimation: a compendium and appraisal. Ecology and Evolution 2:2357–2365.

R Core Team (2021). R: A language and environment for statistical computing. R Foundation for Statistical Computing, Vienna, Austria. URL https://www.R-project.org/.

Sved JA, Feldman MW (1973). Correlation and probability methods for one and two loci. Theor Popul Biol 4: 129–132.

Weir BS, Hill WG (1980). Effect of mating structure on variation in linkage disequilibrium. Genetics 95: 477–488.

Wang, J., 2001. A pseudo-likelihood method for estimating effective population size from temporally spaced samples. Genetics Research, 78(3), pp.243–257.

Wang, J., 2009 A new method for estimating effective population sizes from a single sample of multilocus genotypes. Molecular Ecology 18:2148–2164.

Wang J (2016) A comparison of single-sample estimators of effective population sizes from genetic marker data. Molecular Ecology, 25, 4692–4711.

Wang, J. and Whitlock, M.C., 2003. Estimating effective population size and migration rates from genetic samples over space and time. Genetics, 163(1), pp.429–446.

Waples RK, Larson WA, and Waples RS. 2016. Estimating contemporary effective population size in non-model species using linkage disequilibrium across thousands of loci. Heredity 117:233–240.

Waples, R.S. 1989. A generalized approach for estimating effective population size from temporal changes in allele frequency. Genetics 121:379–391.

Waples, R.S. 2006. A bias correction for estimates of effective population size based on linkage disequilibrium at unlinked gene loci. Conservation Genetics 7:167–184.

Waples, R.S. 2016a. Tiny estimates of the *N_e_*/*N* ratio in marine fishes: Are they real? J Fish Biology 89:2479–2504.

Waples, R.S. 2016b. Making sense of genetic estimates of effective population size. Molecular Ecology 25:4689–4691.

Waples, R.S., and C. Do. 2008. *LDNE*: A program for estimating effective population size from data on linkage disequilibrium. Molecular Ecology Resources 8:753–756.

Waples, R.S., and C. Do. 2010. Linkage disequilibrium estimates of contemporary *N_e_* using highly variable genetic markers: A largely untapped resource for applied conservation and evolution. Evolutionary Applications 3:244–262.

Waples, R.S., and J.R. Faulkner. 2009. Modeling evolutionary processes in small populations: Not as ideal as you think. Molecular Ecology 18:1834–1847.

Waples, R.S., T. Antao, and G. Luikart. 2014. Effects of overlapping generations on linkage disequilibrium estimates of effective population size. Genetics 197:769–780.

Waples, R.S., Waples, R. and Ward, E.J., 2020. Pseudoreplication in genomics-scale datasets. bioRxiv (doi:10.1101/2020.11.12.380410).

